# Synapse specific alterations of autophagy are a hallmark of Danon disease

**DOI:** 10.64898/2026.04.14.718098

**Authors:** Beatrice Terni, Maria Quiles-Pastor, Zoë Reynolds, Kelsey Coppenrath, Nikko-Ideen Shaidani, Pablo Martínez San Segundo, Shawn Adam, Nicolás Riffo-Lepe, Zachary Smith, Marko Horb, Carlos Aizenman, Artur Llobet

## Abstract

Danon disease is a rare disorder caused by mutations in the *LAMP2* gene, which encodes a lysosomal membrane protein key to the endolysosomal pathway and autophagy. Affected individuals show multisystemic alterations that include cardiomyopathy, skeletal muscle weakness, visual deficits and cognitive impairment. Here we establish a knockout *LAMP2* line in *Xenopus tropicalis* that reproduces the characteristic cardiac activity, mobility impairments and vision deficits present in the disease. Damaged mitochondria were abundantly found in skeletal muscle fibers. *LAMP2* mutant *X. tropicalis* detected light with a reduced preference for green wavelengths. Visual deficits were consistent with the finding of damaged mitochondria in the inner segment of rods but not in cones. Differences in autophagic flux were found in presynaptic terminals from photoreceptors and olfactory sensory neurons (OSNs), which establish the first synapse processing vision and olfaction, respectively. In wild-type animals autophagic shapes were observed in OSN terminals but were absent from photoreceptor ribbon synapses. In knockout *LAMP2* tadpoles, autophagic organelles covered 7% of the OSN presynaptic terminal surface, a three-fold increase compared to photoreceptor terminals. These differences suggest that *LAMP2* plays synapse-specific roles that could be an important determinant of the psychiatric manifestations present in Danon disease and support the use of *LAMP2 X. tropicalis* to shed new light on the pathological bases of this lysosomal storage disorder.

## INTRODUCTION

Danon disease (DD) is a rare disorder caused by mutations in the *LAMP2* gene, one of the most abundant proteins in the lysosomal membrane that plays a key role in endolysosomal function and autophagy (Cenacchi et al. 2020; Qiao et al. 2023; Saftig et al. 2001). Mutations follow an X-linked dominant inheritance pattern and affect the autophagy pathway, causing multisystemic manifestations including cardiomyopathy, skeletal muscle weakness, psychiatric alterations, cognitive impairment and visual deficits (Yardeni et al. 2017; López-Sainz et al. 2019). DD hemizygous males develop more severe symptoms than heterozygous females, driving to their premature death at an average age of ∼20 years because of severe heart failure (Boucek, Jirikowic, and Taylor 2011). Primary treatments of DD patients are thus targeted to solve cardiac alterations, for example through heart transplant; however, the effects on the nervous system persist throughout life and are poorly understood.

The only experimental evidence available on DD comes from animal models, for example knockout *LAMP2* mice, which reproduce the main hallmarks of this disorder (Tanaka et al. 2000). These mice show altered lysosomal activity as well as the accumulation of p62-positive aggregates and autophagic vacuoles within hippocampal neurons and their presynaptic terminals (Rothaug et al. 2015). Although these observations confirm the involvement of *LAMP2* in neuronal autophagy, it is necessary to shed light on synapse specific alterations occurring in DD. The endolysosomal system plays a fundamental role in synaptic homeostasis (Spencer, Sudarikova, and Devine 2025), autophagosome biogenesis is linked to the synaptic vesicle cycle (Yang et al. 2022) and autophagy participates in the pruning of dendritic spines (Kallergi et al. 2022), which altogether suggest the synaptic roles of *LAMP2* could be relevant to psychiatric manifestations present in DD patients.

DD has been experimentally modeled in mouse, rats and zebrafish (Saftig et al. 2001; Ma et al. 2018; Dvornikov et al. 2019) and here, we expand the current toolkit by editing through CRISPR/Cas 9 the *Xenopus tropicalis* genome to establish a mutant *LAMP2* line (Tandon et al. 2017). Mutant tadpoles showed histological and functional hallmarks of DD in muscle and nervous tissues, thus reproducing observations obtained in other species. Here, we take advantage of the unique experimental possibilities of *Xenopus* tadpoles to associate synaptic function to behavior (Pratt and Khakhalin 2013) and provide evidence for synapse specific roles of *LAMP2* that could explain the variable psychiatric manifestations present in DD patients.

## MATERIALS AND METHODS

### Generation of *LAMP2* knockout *X. tropicalis*

All animal experiments were done in accordance with the Marine Biological Laboratory Institutional Animal Care and Use Committee (IACUC protocol No 24-33). Seven guide RNAs were designed using CRISPRdirect (https://crispr.dbcls.jp/) (Naito et al. 2015) to target the first exon of *Xenopus tropicalis LAMP2*. Each guide received 5’ GG dinucleotide modifications to increase mutagenetic activity (Gagnon et al. 2014). T7 MEGAscript kit (Ambion, Cat. No. AM1334) was used to synthesize sgRNAs. *X. tropicalis* wild-type Nigerian (RRID: NXR_1018) animals were used to generate F0 founders. Embryos at 1-cell stage were microinjected into the animal region with 500 pg of sgRNA and 1000 pg cas9. Of the seven guides tested, only sgRNA5 (GGATGGCAAAGCACCTATAC) yielded any cutting. DNeasy Blood & Tissue Kit (Qiagen, Cat. No. 69506) was used for the isolation of genomic DNA. Forward primer 5’- CTAATACCCTACCTTTTAACCC - 3’ and reverse primer 5’- TACAGACATCATTGCAGTTAGC - 3’ were used for PCR amplification and samples were cleaned using Nucleospin PCR Clean-up procedure (Macherey-Nagel 740609.250). All mutations were confirmed by sanger sequencing. Tissue samples were collected from adult frogs from the webbing on their hindlimbs. Once founders reached sexual maturity, animals were outcrossed to wild types to test for germline transmission at the F1 generation. Two of these F0 females produced favorable mutations that resulted in seven total adult F1 animals with heterozygous mutations (one -2/+, two +5/+, two -5/+, and two - 13/+ animals) and seven adult F1 wild type animals. Intercrosses of the -13 bp mutation and - 2 bp mutation lines were conducted to generate mutant F2 *LAMP2* tadpoles used for this phenotypic characterization. The -13 bp mutation results in a frameshift at amino acid 80 and an early stop codon induced ten amino acids downstream while the -2 bp deletion causes a frameshift at amino acid 81 and an early stop codon induced three amino acids downstream.

### Line Accessibility

Both the -13 bp mutation (Xtr.*LAMP2*emHorb; RRID: NXR_3182) and -2 bp mutation (Xtr.*LAMP2*em2Horb; RRID: NXR_3219) lines were used for this study and are available at the National Xenopus Resource (NXR) (http://mbl.edu/xenopus/, RRID:SCR_013731).

### Husbandry

Wild type *X. tropicalis* (RRID: NXR_1018) animals used for generating the *LAMP2* mutants were housed at the National Xenopus Resource (NXR) following established protocols (McNamara, Wlizla, and Horb 2018; Shaidani et al. 2020). *LAMP2* tadpoles used in this study were generated by intercrossing two -13 bp mutants or by crossing one with a -2 bp mutant through natural fertilization. To induce egg laying, females were given 20 U of pregnant mare serum gonadotropin (PMSG; BioVender, RP17827210000) and 200 U of human chronic gonadotropin (hCG; BioVender, RP17825010), while males received 10U of PMSG and 100U of hCG to induce mating behavior (McNamara, Wlizla, and Horb 2018). Tadpoles were obtained by natural mating and kept in water tanks at 25 °C., were grown in 0.1 x Marc’s Modified Ringers (MMR) solution and were used for the experiments at stages 47-50 of the Nieuwkoop–Faber criteria (Nieuwkoop and Faber 1956).

### Histology

Tadpoles at NF stages 50-52 were anesthetized in 0.02 MS-222 and fixed at 4°C in a 4% paraformaldehyde solution prepared in PBS or 1.2 % glutaraldehyde solution prepared in 0.05M PBS to process them for optical or electron microscopy, respectively. Distal tail fragments were transected for genotyping before fixation. For optical microscopy tadpoles were embedded in paraffin and sectioned at 12 μm thickness. Staining was carried out using hematoxylin-eosin or immunohistochemistry following the EnVision + system peroxidase procedure (Dako, Denmark). For immunohistochemistry sections were incubated with one of the primary antibodies: LAMP2 mouse monoclonal antibody (Cat. No., 66301, Proteintech) or LAMP1 rabbit polyclonal antibody (Cat. No., ab24170, Abcam) at a dilution of 1:100. HRP conjugated secondary antibodies were incubated at a 1:500 dilution. The peroxidase reaction was visualized with diaminobenzidine and H_2_O_2_. For electron microscopy, animals were post fixed in 1% osmium tetroxide/1.5% potassium ferricyanide in 0.1M of sodium cacodylate buffer, dehydrated in alcohol series and embedded in Epon resin. 1µm thick sections were stained with toluidine blue for the identification of regions of interest. Ultrathin sections (60 nm) were contrasted with uranyl acetate and lead citrate and viewed under a JEOL 1010 electron microscope. Random sections obtained in three WT and three *LAMP2 ^(- /-)^* tadpoles were used for analysis. The area covered by autophagic intermediates and endolysosomal organelles was divided by the presynaptic terminal area in images obtained at 60K or 80K magnification.

### Tadpole mobility assays

Tadpoles at NF stage ∼50 were placed in a 100 mm diameter dish containing 0.1X MMR and were allowed to settle for 2 minutes. Movies of 1 min duration were acquired at 30 Hz using a camera positioned 20 cm over the dish. Image sequences were opened in Image J. After subtracting the maximum intensity projection from each frame, tadpole positions were tracked using the wrMTrck plugin (Nussbaum-Krammer et al. 2015). Data were imported in Igor Pro 9 (Wavemetrics, OR) to quantify the average swimming velocity, the maximum swimming velocity and quiescent periods for each tadpole.

### Heart Function Analysis

In order to assess heart function, tadpoles were anesthetized with 0.02% MS-222 by immersion. Within 2 minutes after swimming cessation, they were placed ventral side up under a stereomicroscope and immediately imaged. Because *X. tropicalis* tadpoles are transparent, the heart was imaged at 60 FPS for 30 seconds, taking care that both the ventricle and atria were clearly visible. Videos were imported into FIJI and converted into grayscale TIFF stacks. ROIs were placed on the ventricle and atria (V and A) for analysis. To measure the heartbeat strength, we imaged the amount of blood entering and leaving each chamber during each heartbeat, as measured by the average change in pixel intensity across the ROI (Bartlett et al. 2004), and expressed as the normalized change in intensity from peak to trough. Amplitude of the peaks as well as timing was calculated using Axograph. The Amplitude CV is the coeffcient of variation (CV) of the heartbeat amplitude over the length of the recording, and the V-A time difference is the peak-to-peak timing between the two chambers.

### Electrophysiology

For *in vivo* recordings, animals were anesthetized with 0.02% MS-222 for 10 min and subsequently immobilized by pretreatment with 0.1 mM bungarotoxin for 4 min(Khakhalin et al. 2014). The ACSF bathing solution contained (in mM): 115 NaCl, 4 KCl, 3 CaClL, 3 MgClL, 5 HEPES, and 10 glucose, adjusted to pH 7.2, 255 mOsm/kg (Thompson and Aizenman 2024). Animals were secured onto a Sylgard block shaped to stabilize the preparation, using three dissecting pins and an overhead staple(Xu et al. 2011). To record from the optic tectum (OT), the tectal lobes were exposed by removing the skin over the right tectum and severing both the caudal and rostral dorsal commissures connecting the two tecta. OT neurons were visualized with a light microscope equipped with a 60× water-immersion objective and an infrared CCD camera (Scientifica Slice Scope Pro 6000).

Loose cell-attached recordings and whole-cell voltage-clamp recordings were conducted at room temperature (22–25 °C) using glass micropipettes with resistances of 9–12 MΩ. Pipettes were filled either with ACSF (for loose cell-attached recordings) or intracellular solution containing (in mM): 100 K-gluconate, 8 KCl, 5 NaCl, 1.5 MgCl2, 20 HEPES, 10 EGTA, 2 ATP disodium salt hydrate, 0.3 GTP sodium salt hydrate, pH 7.2, 255 mOsm/kg). Cells were sampled randomly from the central half of the tectum.

For loose cell-attached recordings, cells were approached in current-clamp configuration, and gentle negative pressure was applied until clear spike activity could be distinguished from background noise. These recordings were defined by seal resistances between 40 and 200 MΩ. Cells that failed to generate spikes in response to changes in illumination were excluded from recording. A fiber optic cable positioned in front of the eye delivered a 1 s full-field visual stimulus produced by a white LED. Light intensity was calibrated to produce three stimulus levels corresponding to 50%, 75%, and 100% of the maximal stimulus intensity, where the maximal intensity was defined as the level that elicited the peak response. ON responses were quantified.

Whole-cell patch-clamp recordings were performed at holding potentials of either −45 mV or +5 mV (junction potential not corrected) to isolate excitatory and inhibitory synaptic currents, respectively. Electrophysiological signals were amplified with a Multiclamp 700B amplifier (Molecular Devices, Union City, CA, USA), digitized at 10 kHz using a Digidata 1550 interface (Axon Instruments), and acquired with pClamp 11 software. Subsequent data processing and analysis were carried out offline using Clampfit, part of the pClamp suite. The whole-brain preparation was carried out following the procedure described by Wu et al. (Wu, Malinow, and Cline 1996). Animals were first anesthetized using 0.02% MS-222 prepared in 10% Steinberg’s solution. The dorsal skin was removed to expose the brain, and a dorsal midline incision was made from the base of the spinal cord to the olfactory bulbs. The brain was then carefully dissected and transferred to a recording chamber containing room-temperature HEPES-buffered extracellular saline (115 mM NaCl, 4 mM KCl, 3 mM CaClL, 3 mM MgClL, 5 mM HEPES, and 10 mM glucose; pH 7.2, 255 mOsm/kg). The tissue was positioned on a Sylgard block with the ventricular surfaces oriented upward. Stabilization was achieved by inserting shortened insect pins through the caudal hindbrain and through one or both olfactory bulbs. To stimulate retinal ganglion cell axons, a bipolar stimulating electrode composed of two adjacent 25-µm platinum leads (CE2C75; FHC, Bowdoin, ME) was positioned at the optic chiasm (OC).

### Color and light preference assay

We adapted the open-field color preference assay originally developed for *X. laevis* (Hunt, Bruno, and Pratt 2020) for use with *X. tropicalis*. Because *X. tropicalis* tadpoles are smaller than *X. laevis* at equivalent developmental stages, we used a 10-cm-diameter circular Petri dish filled with Steinberg’s solution to a depth of 10 mm. The dish was placed on a 7-inch IPS LCD touchscreen (Hosyond; 1024×600 pixels; HDMI interface) oriented face-up to project stimuli onto the dish floor. The outer edge of the dish was wrapped with black tape to occlude extraneous visual stimuli. The entire apparatus was enclosed in a custom 3D-printed light-tight enclosure so that the LCD monitor provided the sole source of illumination. Ten NF stage 49 tadpoles were randomly selected and placed in the dish for each trial; a fresh group was used for each condition.

### Visual stimuli

Visual stimuli were generated and presented using Psychtoolbox-3 running in MATLAB (The MathWorks). The display was divided into four equal quadrants. For the light-preference assay stimuli consisted of a black background with one quadrant in white. Tadpoles normally swim towards the light quadrant. After one minute an image was taken and tadpoles on the light quadrant were counted and a new quadrant was randomly illuminated. Stimuli were presented 30 times. For the color preference assay, one quadrant was filled with a colored stimulus; the remaining three quadrants displayed a uniform background. We tested three stimulus colors: green (RGB 0/216/0), blue (RGB 0/33/221), and red (RGB 232/20/42). These RGB values were calibrated to equate luminance across hues using the display’s gamma function in Psychtoolbox-3. Two background conditions were tested: white (RGB 255/255/255) and black (RGB 0/0/0). The full factorial combination of 3 colors × 2 backgrounds yielded six stimulus conditions per genotype, results for a given color were combined. Each trial began with a 60-s acclimation period during which the monitor displayed a uniform grey field (RGB 128/128/128). The grey field was then replaced by the color-versus-background stimulus for 30 min. At the end of the trial the display returned to grey. Stimulus onset, duration, and identity were logged automatically.

### Video acquisition and tracking

We recorded tadpole behavior from above using a Raspberry Pi Camera Module 3 (Raspberry Pi Ltd) mounted approximately 12 cm above the dish. Video was captured at 30 frames per second at 1080p resolution (1920×1080 pixels) for the full 30-min trial duration. All trials were conducted in the morning (before 12:00) to minimize circadian variation in visual behavior. Behavioral testing was conducted at room temperature (22–23°C). For the color preference assay, we extracted individual tadpole positions from video recordings using SLEAP (Social LEAP Estimates Animal Poses, v1.4.1a2)(Pereira et al. 2022). We used a top-down approach in which a detection network first localized individual animals and a pose-estimation network then predicted the body centroid of each tadpole within each detected crop. Predicted centroid tracks were proofread in the SLEAP graphical interface to correct identity swaps and missed detections.

### Data analysis and statistics

Tracked centroid coordinates were exported from SLEAP as HDF5 analysis files and processed using custom MATLAB scripts. To quantify spatial preference, we defined the color-patch quadrant as a polygonal region of interest (ROI) occupying 25% of the arena and performed a frame-by-frame point-in-polygon test on each centroid trajectory. For each tadpole we computed the dwell fraction: the number of frames the centroid fell within the ROI divided by the total number of visible (tracked) frames. Under the null hypothesis of no spatial preference, the expected dwell fraction is 0.25. Statistical analysis and graphing were performed in GraphPad Prism. Unpaired student’s t-test was used to compare two groups and ANOVA analysis was applied when comparing three or more groups. Data are expressed as mean±s.e.m.

### In vivo imaging of endolysosomal organelles

The two nasal cavities of tadpoles anesthetized with 0.02% MS-222 were filled with 0.15–0.3 µl of 1mM lysotracker deep red (Thermofisher Scientific, Walthman, MA, Reference: L12492). A microinjector delivered a puff of the dye (30 psi, 6 ms) using pipettes of ∼1 μm tip diameter fabricated from borosilicate glass capillaries. Animals recovered from anesthesia in 2 L tanks filled with tadpole water. Imaging sessions started by anesthetizing tadpoles in water containing 0.02% MS-222 to remove the skin covering the olfactory bulb. The goal was to eliminate melanocytes that could interfere with imaging. Anesthetized animals were placed dorsally on a 25 mm diameter coverslip and embedded in a 1% solution of low melting point agarose (Sigma, Sant Louis, MO, Reference: A9414-10G). Imaging experiments were carried out 18-24 h after application of lysotracker deep red in a Zeiss LSM 880 confocal microscope (Carl Zeiss AG, Oberkochen, Germany). An imaging session lasted 10-15 min. Confocal sections were acquired using a 63X glicerol immersion objective LD LCI Plan-Apochromat (1.2 N.A.). Lysotracker deep red was excited at 633 nm and fluorescence was collected at 700 nm. Tracking of labelled acidic organelles was carried out in 1024x1024 pixel confocal sections at a resolution of 7.59 pixels/μm^-1^ (voxel size 0.132x0.132x0.499 μm^3^). A time series of a single 2D confocal section was collected at 0.8 Hz for 4 minutes. The selected region was situated dorsally to maximize light transmission.

Image analysis was carried out using Image J and Fiji. The drift in the X-Y plane was corrected using the StackReg plugin(Thévenaz, Ruttimann, and Unser 1998). MosaicSuite plugins (Sbalzarini and Koumoutsakos 2005) were used to select image trajectories, as well as the diffusion coefficient of identified acidic organelles. Only trajectories of single particles that were tracked for >45 s were considered. Particle positions were imported in Igor Pro 9 (Wavemetrics, Lake Oswego, OR) to quantify mobility.

## RESULTS

### *LAMP2* mutant *X. tropicalis* tadpoles show characteristic alterations of Danon Disease in muscle tissue

Adult *X. tropicalis* carrying heterozygous mutations of *LAMP2* ^(−2/+,-13/+)^ were intercrossed to generate mutant (^+/-^ and ^-/-^) tadpoles (Fig. 1A). Growth of WT and mutant larvae was comparable and occurred through a comparable time course. No obvious differences were observed in any external phenotype as tadpoles progressed through developmental NF stages (Nieuwkoop and Faber 1956). Gross histological analysis did not show alterations in organ size (Fig. 1B), however, the inspection of tail muscle fibers in *LAMP2 ^(-/-)^* tadpoles evidenced some abnormalities. The characteristic pattern of light and dark bands was not obvious and there were large extracellular spaces among fibers (Fig. 1C). These alterations were attributed to the absence of functional *LAMP2* as revealed by immunostainings. The antibody recognizing a peptidic sequence found between residues 235-339 detected *LAMP2* in WT tadpoles (Fig. 1D) but did not provide any signal in *LAMP2 ^(-/-)^* tadpoles (Fig. 1E). This finding was consistent with the generation of a truncated *LAMP2* isoform caused by the presence of an early stop codon (Fig. 1A) that was likely nonfunctional (Qiao et al. 2023). Endolysosomes were still present in muscle fibers of mutant *LAMP2* tadpoles because the distribution of LAMP1, another major membrane lysosomal protein, was not affected by the absence of *LAMP2* (Figs. 1F and G).

**Figure 1.**
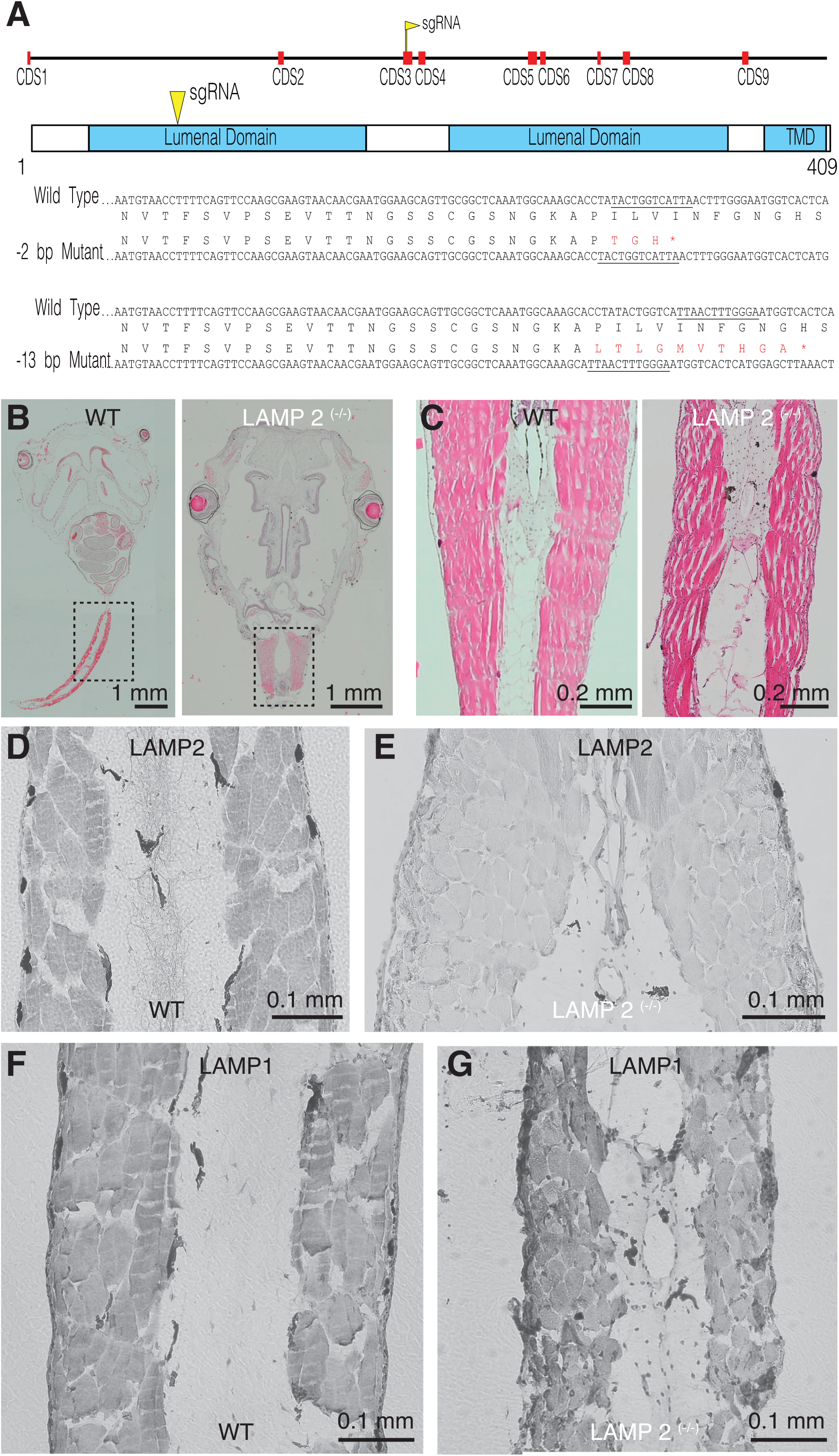
Generation of knockout *LAMP2 X. tropicalis*. **A)** sgRNAs targeting a region codifying for the first lumenal domain of *LAMP2* were injected in F0 embryos. F1 animals carrying heterozygous mutations (−2/+ and -13/+) were intercrossed to generate mutant F2 *LAMP2* tadpoles for phenotypic characterization. The -13 bp mutation results in a frameshift at amino acid 80 and an early stop codon induced ten amino acids downstream while the -2 bp deletion causes a frameshift at amino acid 81 and an early stop codon induced three amino acids downstream. **B)** The morphology of *LAMP2* ^(-/-)^ tadpoles was comparable to WT animals, revealed by sagittal sections stained with hematoxylin-eosin. **C)** Skeletal muscle fibers present in the tail of *LAMP2* ^(-/-)^ tadpoles showed large intercellular spaces. D, E) Immunohistochemistry of tail skeletal muscle fibers against *LAMP2* confirmed the absence of the protein in *LAMP2* ^(-/-)^ tadpoles. The expression of LAMP1 reamined unaltered.

Optical microscopy observations were confirmed by electron microscopy. The characteristic appearance of skeletal muscle fibers found in WT tadpoles (Figs. 2A and B) was not observed in *LAMP2 ^(-/-)^* animals. Skeletal muscle fibers showed irregular sizes and large intercellular spaces filled with vacuoles (Figs. 2C and D). Two evident intracellular alterations in fibers of *LAMP2 ^(-/-)^* tadpoles were a swollen sarcoplasmic reticulum and mitochondria containing fragmented cristae. These morphological alterations were correlated to mobility deficits. WT tadpoles swim by alternating active and idle periods, which resulted in an average velocity of ∼5 mm/s (Figs. 2E and F). Quiescent periods were more frequent and longer in mutant *LAMP2* tadpoles (^+/-^ and ^-/-^) so that they only moved during 15-20% of the observation time. The consequence was a decrease in velocity to ∼2.5 mm/s. No differences in swimming behavior were observed between *LAMP2* (^+/-^) and *^(-/-)^* tadpoles. Interestingly, the only parameter analyzed that was not significantly different between WT and mutant *LAMP2* tadpoles was maximum swimming velocity. Taken together these results evidenced the presence of a skeletal myopathy, a key characteristic of DD.

**Figure 2.**
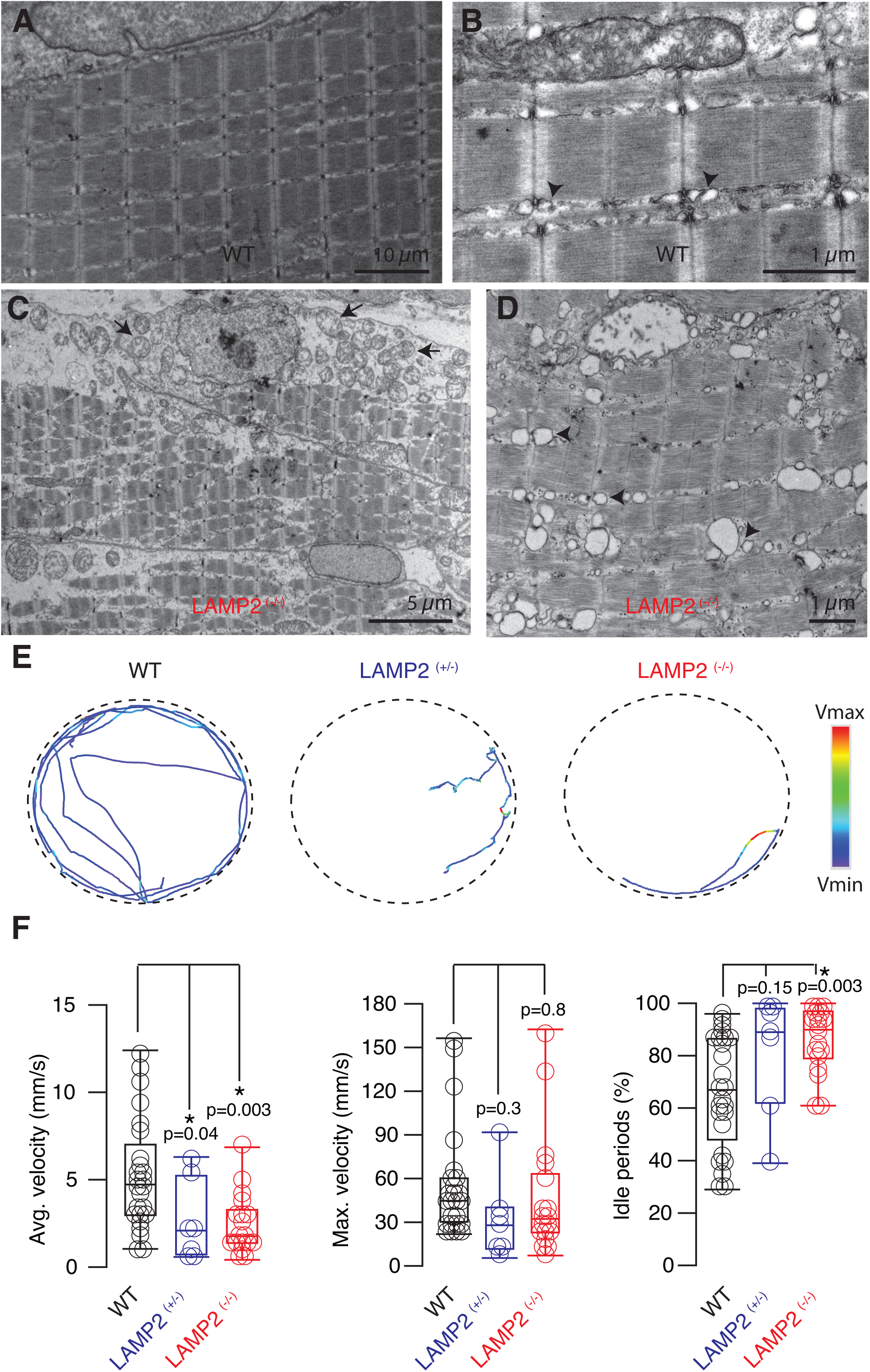
*LAMP2* ^(-/-)^ tadpoles show deficiencies in swimming associated to alterations in skeletal muscle fibers. **A-D**) The morphology of skeletal muscle fibers (A,B) was altered in *LAMP2* ^(-/-)^ tadpoles (C,D). Mitochondria were electrolucent and the organization of cristae was disrupted (C, arrows). Cisternae of the sarcoplasmic reticulum were enlarged, driving to a disorganization of triads (B,D, arrowheads). **E)** The mobility of single tadpoles was tracked over 60 s in a 100 mm diameter petri dish. Swimming of WT animal was characterized by an alternation of forward movements with quiescent periods. Mutant *LAMP2* animals moved aberrantly, displaying frequent body bends that generated a circular swimming as well as long quiescent periods. F) Box plots showing the median (horizontal line), 25 to 75% quartiles (boxes), and ranges (whiskers) of the average velocity, maximum velocity and quiescent periods found for individual tadpoles (circles) in the three experimental groups studied: WT, *LAMP2* ^(+/-)^ and *LAMP2* ^(-/-)^.

Further confirmation of abnormalities in muscle tissue was obtained by studying cardiac activity in *LAMP2 mutants* (^+/-^ and ^-/-^; Fig. 3). Tadpoles were anesthetized with MS-222 by immersion and would stop swimming within 1-2 minutes. Within 2 minutes after swimming cessation, they were placed ventral side up under a stereomicroscope. Because *X. tropicalis* tadpoles are transparent, the heart was imaged at 60 FPS for 30 seconds, taking care that both the ventricle and atria were clearly visible, and ROIs were placed on the ventricle and one of the atria for analysis (Figs 3A and B; see methods). The strength of the heartbeat could be measured by measuring the amount of blood entering and leaving each chamber during each heartbeat, measured as a change in pixel intensity across the ROI, and calculating the normalized change from peak to trough. We found that *LAMP2 mutant* tadpoles had significantly lower heartbeat amplitudes than WT (Figs 3A and C, WT: 4.9±1.3%, n=9; *LAMP2 mutants*: 1.7±0.29%, n=11; Student’s t-test, p=0.017), as well as more beat-to-beat variability, as indicated by the CV of the amplitude (WT: 0.37±0.06; *LAMP2 mutants*: 0.54±0.04; unpaired Student’s t-test p=0.037). When we looked at timing, we found that there was no significant difference in heart rate (WT: 125±8 BPM; *LAMP2 mutants*: 137±5 BPM; p=0.245). We did find a small but significant difference between the time difference between peak contractions between atria and ventricles (C: 241±23 ms; *LAMP2 mutants*: 179±19 ms; p=0.038, unpaired Student’s t-test). No differences were found between homozygotic ^(-/-)^ and heterozygotic mutants ^(-/+)^. While the heart rhythm seems largely unaffected, the integrity of the heart muscle might be compromised in the *LAMP2* mutants, as indicated by weaker contractions. Altogether, the data obtained in skeletal and cardiac muscle (Figs. 2 and 3) illustrate that *X. tropicalis LAMP2* mutant tadpoles faithfully reproduce key features of DD.

**Figure 3.**
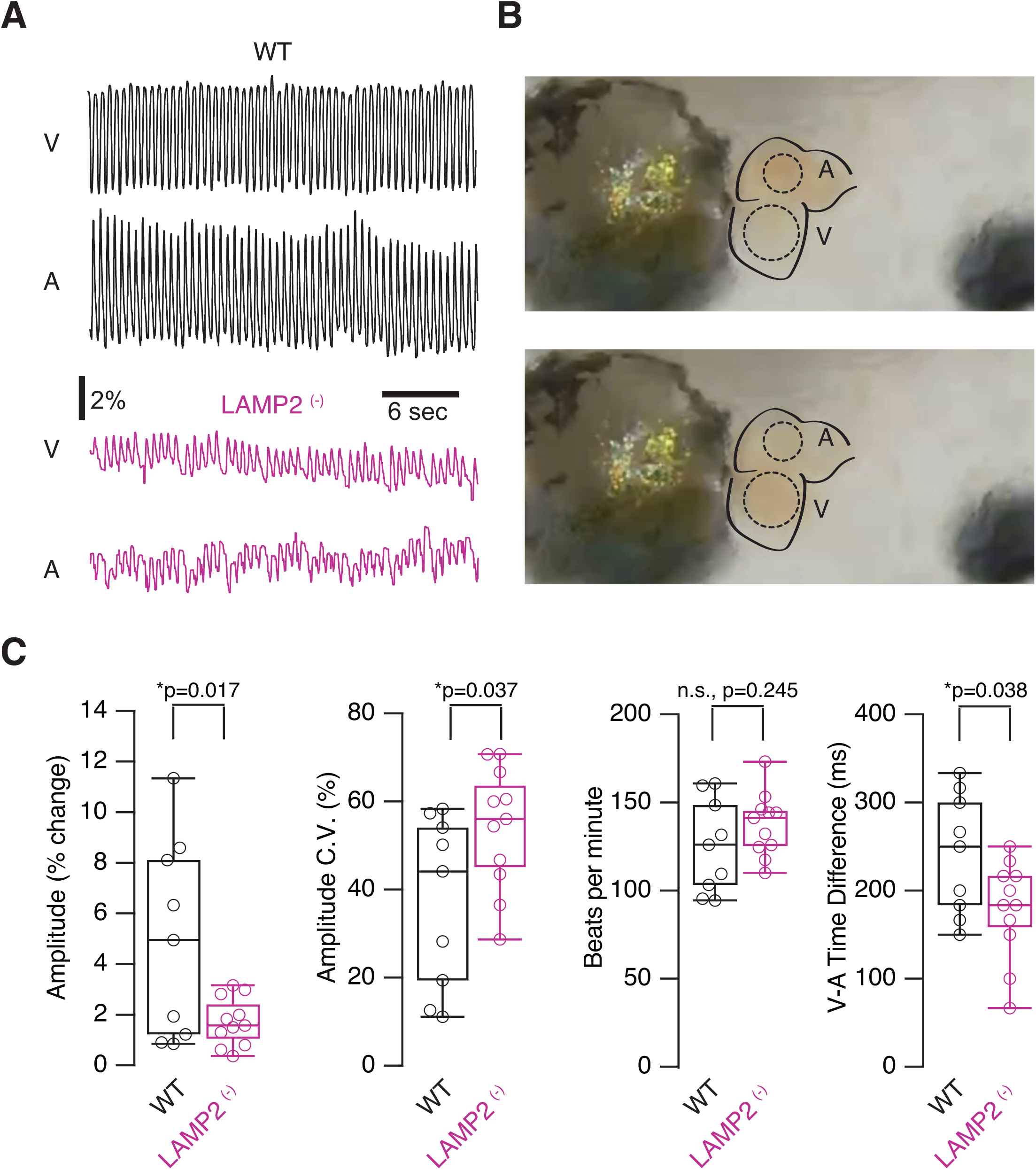
LAMP2 ^(-/-)^ tadpoles show altered heart function. **A)** Sample normalized traces of heartbeat function from ventricle and atrial chambers for control and mutant tadpoles. Oscillations represent changes in grayscale value measured in each heart chamber over a 30 second period. Notice the shallower oscillations in the mutants. **B)** Sample heart image showing ROIs over the atrium and ventricle. **C)** Comparison of various measures of heart function including heartbeat ejection amplitude, beat to beat CV of the amplitude, beats per minute and timing between V and A peaks. * p<0.05, Student’s t-test.

### Visual deficits and retinal alterations are present in *LAMP2* mutant *X. tropicalis* tadpoles

Visual deficits are commonly present in DD patients and are associated to a thinning of the photoreceptor layer as well as to the disruption of retinal pigment epithelium functionality (Chaudhary and Singh 2025). Histological observation of the retina in *LAMP2 ^(-/-)^* tadpoles evidenced a non-homogenous thickness of the photoreceptor layer and an altered external plexiform layer (Figs. 4A and B). Both observations suggested that photoreceptor integrity was compromised. In WT tadpoles, the inner segment of rods and cones is populated by mitochondria showing numerous cristae and an electrodense matrix (Figs. 4D and E). Mitochondria were also abundant in the inner segment of *LAMP2 ^(-/-)^* tadpole photoreceptors; however, those found in rods were electrolucent (Figs. 4F and G). The cause was a decrease in the density and length of cristae. The disruption of mitophagy could be a possible reason, as described in pluripotent stem cell-derived cardiomyocytes from DD patients (Hashem et al. 2017); however, the effects were cell-specific. Mitochondrial alterations visualized in rods resembled those present in skeletal muscle fibers (compare Figs. 2D and 4G) but they were not observed in cones of *LAMP2 ^(-/-)^* tadpoles (Fig. 4F). The presence of different alterations to mitochondria indicates that the role played by *LAMP2* must be interpreted in a cell-autonomous context.

**Figure 4.**
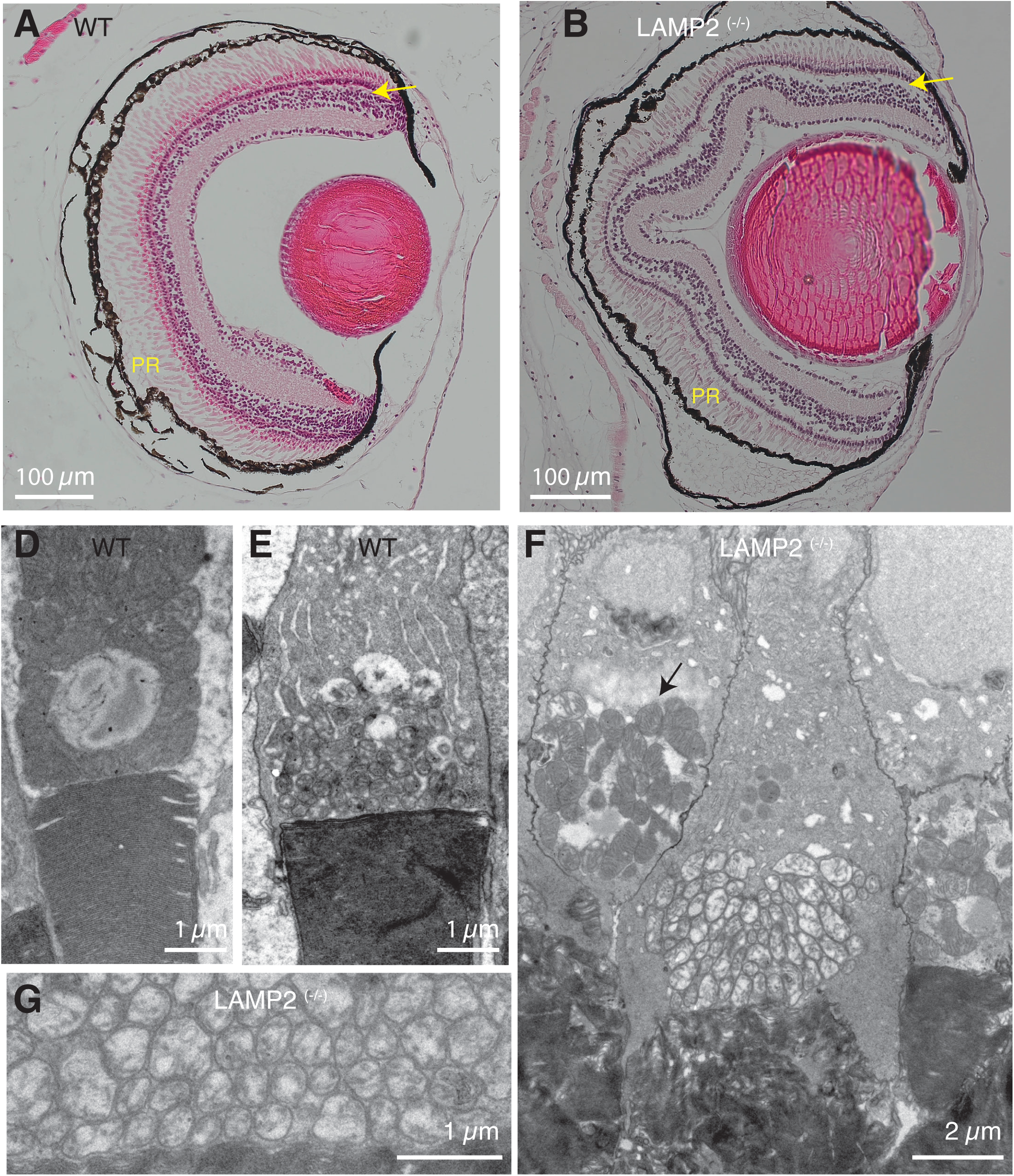
*LAMP2* ^(-/-)^ tadpoles show retinal alterations. **A,B**) Eye sagittal sections suggesting a decreased photoreceptor (PR) density in *LAMP2* ^(-/-)^ tadpoles and a disruption of the outer plexiform layer (arrow). **D,E)** Abundant mitochondria are found in the inner segment of cones (D) and rods (E ) of WT tadpoles. **G, F)** Rod mitochondria appear electrolucent and the organization of cristae is disrupted in *LAMP2* ^(-/-)^ tadpoles (G). In contrast, cone mitochondria (arrow) do not show obvious alterations (F).

To correlate the morphological observations of the retina to visual function, we began by testing *LAMP2* mutants in a simple light/dark place preference assay. *X. laevis* tadpoles are known to prefer a light background over a black one (Bruno et al. 2022). Firstly, we assessed whether this preference was conserved in WT *X. tropicalis* tadpoles, and secondly, whether this preference was seen in the *LAMP2* mutants. Tadpoles were placed in groups of 10 in a transparent round chamber over an LED screen in which one random quadrant was white and the rest of the arena was black (Fig. 5A). Every minute over a course of 30 minutes, an image was taken, and the number of tadpoles in the white quadrant was recorded (Fig .5B). We found a significant preference for the white quadrant for both WT and *LAMP2* mutants (^+/-^ and ^-/-^, Fig 5C; WT: 6.06±0.47 tadpoles, p=0.016;, *LAMP2 mutants*: 5.19±0.75 tadpoles, p=0.023, one-sample T-test), indicating that despite observed ultrastructural deficits seen in the retina, basic visual function was preserved.

**Figure 5.**
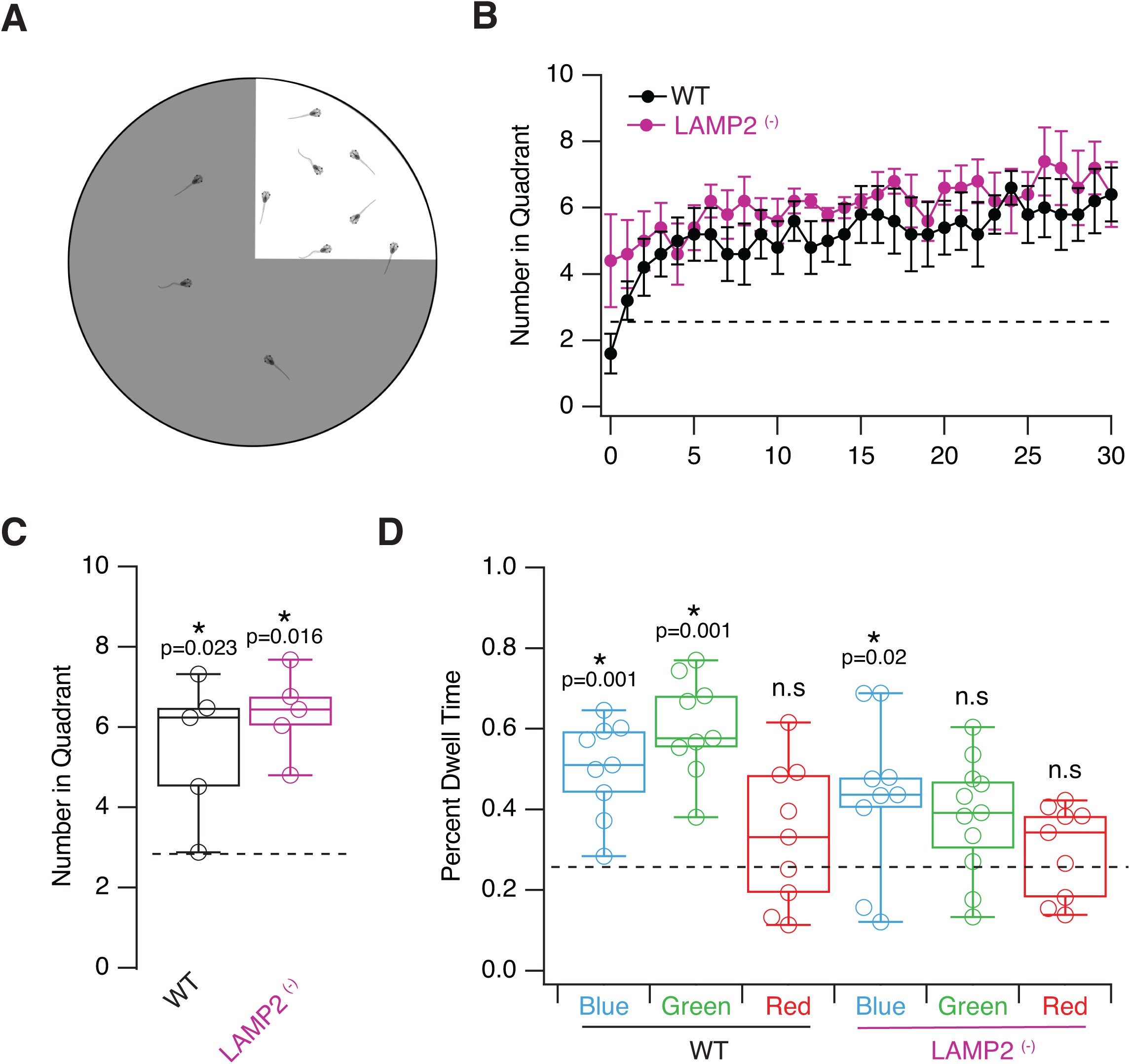
Visual behavioral differences in LAMP2 ^(-)^ tadpoles. **A)** Sample layout of the quadrant preference test, where tadpoles are allowed to swim freely and time spent on the stimulus-presenting quadrant. **B)** Tadpoles present in light quadrant (vs dark) monitored every minute over a 30 minute period indicate a preference for light over dark in both WT and LAMP2 (-) tadpoles. Tadpoles were tested in groups of 10 with 5 runs for each condition. **C)** Average number of tadpoles over the 30 minute period show significant preference for a light over dark area for both groups. **D)** Color preference tests measuring percent dwell time of individual tadpoles for backgrounds of different colors. WT tadpoles showed significant preference for blue and green background, but not for red. LAMP2 (-) tadpoles only showed a preference for blue backgrounds but not green, indicating rod dysfunction. *** >0.001, ** >0.01, * >0.05, one sample t-test (see methods).

We next tested whether color vision was affected. A place preference assay where a single quadrant was presented alternately as blue, green or red, and individual tadpoles were tracked to allow us to calculate the dwell time in the colored quadrant. Like *X. laevis* tadpoles (Hunt, Bruno, and Pratt 2020), WT *X. tropicalis* showed a clear preference for the colored quadrant when it was either blue or green, but no preference when the quadrant was red (Fig. 5D, proportion of time in Blue: 0.5±0.11, Green: 0.58±0.13, Red: 0.32±0.17, one-sample T-test p=0.001, 0.001, 0.20, respectively). In *X. laevis* most photoreceptors are rods and 97-98% of them are sensitive to green light (Röhlich and Szél 2000). Only a minority of rods sense blue light and cones detect red light. On these bases, our visual tests support that color vision of *X. tropicalis* and *X. laevis* tadpoles is comparable. *LAMP2* mutants (^+/-^ and ^-/-^) showed a smaller but still significant preference to blue and showed no preference for green, and like WT, they did not prefer red light (proportion of time, Blue: 0.43±0.065, Green :0.35±0.17, Red: 0.29±0.03, one-sample T-test p=0.02, 0.06, 0.27, respectively). These data are consistent with the presence of alterations in rods (Fig. 4) and are aligned with the report of a cone-rod dystrophy in DD patients (Thiadens et al. 2012).

### Retinal dysfunctions affect the processing of visual information by tectal neurons

In order to directly test visual responses in the visual areas of the brain, we performed in vivo recordings from the optic tectum. NF stage 49 tadpoles were treated with 0.1 mM alpha-Bungarotoxin and then placed in a recording chamber with extracellular recording solution. A fiber optic cable was placed in front of the eye to generate a 1 second whole field visual stimulus driven by a white LED. Cell attached recordings were performed with a micropipette filled with external medium, which allowed us to record single spikes (Fig. 6A). Light intensity was calibrated to provide three stimulus intensities 50%, 75% and 100% of the maximal stimulus intensity, the maximal intensity being the one that generated a peak response. ON responses were measured. We compared WT controls with both heterozygotic and homozygotic *LAMP2 mutants* (Fig 6B). Since no differences were found between *^(-/-)^*and (^+/-^) *LAMP2* tadpoles, the information of both groups were combined. Across intensities we found that in the *LAMP2 mutants* the ON visual response was significantly weaker (Fig 6C, p=0.004, 2-way ANOVA), and the onset response longer than the WT (Fig 6C, p=0.004, 2-way ANOVA). This indicates that while the *LAMP2* mutants could generate visual responses, they were significantly impaired compared to WT. The optic tectum generates a substantial amount of spontaneous activity. We found that *LAMP2* mutants showed significantly decreased spontaneous activity as evidenced by longer inter-spike intervals and lower spontaneous spike frequency (Fig 6D, WT: 1.02±0.37 Hz, *LAMP2 mutants*: 0.14±0.03 Hz, p<0.001 Mann-Whitney test).

**Figure 6.**
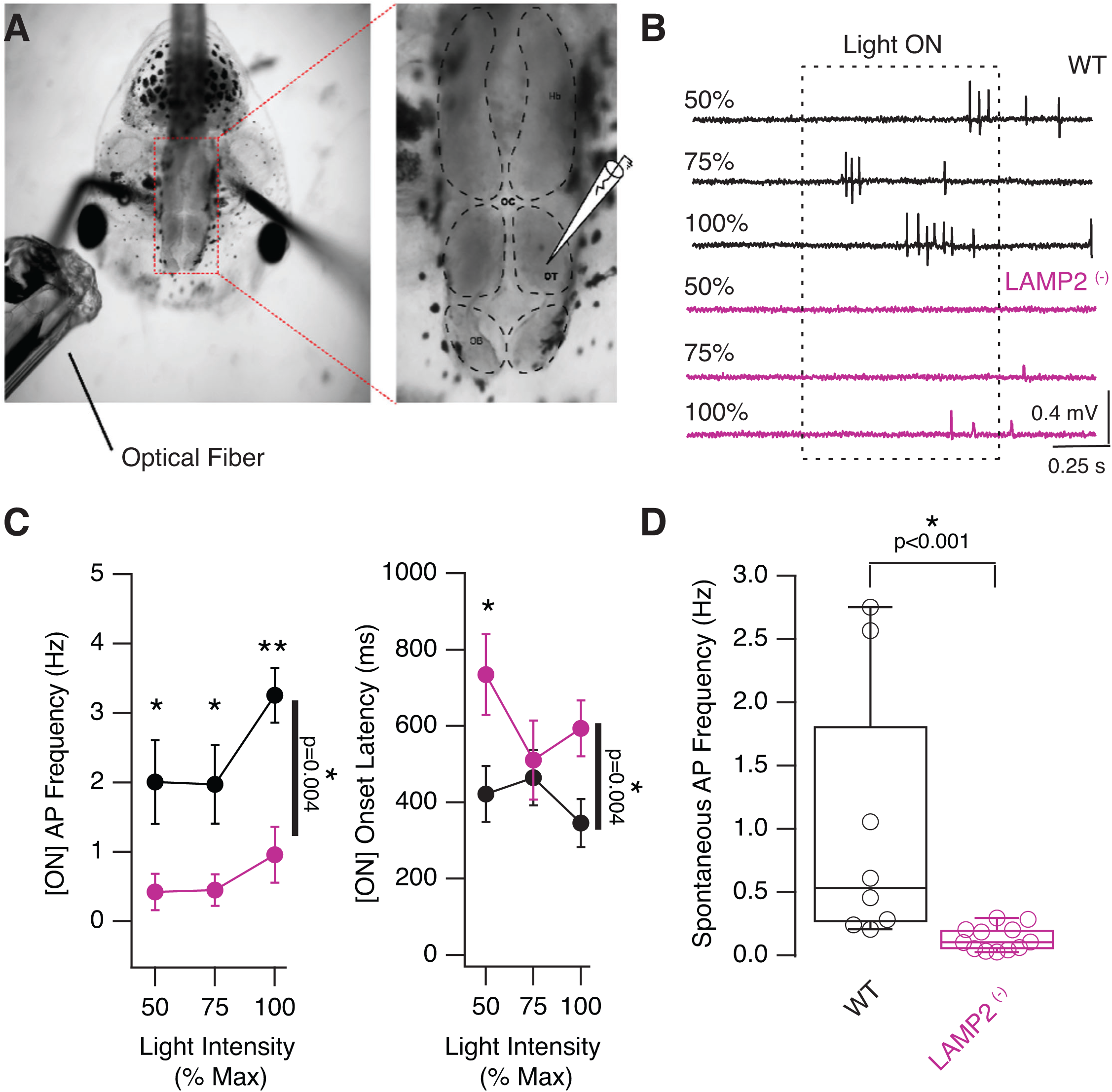
Decreased evoked visual responses are reduced in *LAMP2* ^(-)^ mutants. **A)** *In vivo* electrophysiological recording configuration, shows placement of fiber optic stimulator next to right eye to deliver a whole field visual stimulus. Image on the right shows placement of extracellular recording electrode in the optic tectum. **B)** Sample evokes visual responses using cell-attached recordings at a range of stimulating intensities. Notice the decreased responsiveness in the mutant tadpole. **C)** Average response frequency and onset latency in optic tectal neurons in response to increasing stimulus intensity in control (n=8) and mutant (n=12) tadpoles, shows significantly decreased responsiveness in LAMP2 mutant tadpoles. **D)** Tectal neurons in LAMP2 mutants show very little spontaneous spiking activity, in contrast to WT tadpoles.

To test whether the decrease in visual responses could be due to a dysfunction in retinotectal synaptic transmission, we did electrophysiological recordings in ex vivo whole-brain preparations (Wu, Malinow, and Cline 1996) where the entire brain was dissected and placed in a recording chamber. A bipolar extracellular stimulating electrode was placed in the optic chiasm to directly shock the optic nerve, and whole-cell intracellular recordings were made in tectal neurons. Short-term synaptic plasticity was assessed using paired stimuli (Suppl. Fig. 1A). Paired pulse facilitation (PPF), which is inverse to presynaptic release probability was not different between WT and *LAMP2* mutants (Fig S1A; WT: 1.66±0.16, n=10; *LAMP2 mutants*: unpaired Student’s t-test, 1.46±0.2, n=9; p=0.46).

Most synaptic transmission in the optic tectum of *X. tropicalis* relies on axodendritic contacts formed by presynaptic boutons of sensory neurons that contact tectal neurons (Suppl. Fig. 1B). Typically, a presynaptic terminal contained 2.3±0.3 active zones/μm^2^ and a density of 126±15 synaptic vesicles/μm^2^. In WT tadpoles, tectal synapses were virtually devoid of autophagic intermediates but, in contrast, numerous autophagophores and autophagosomes were evident in *LAMP2 ^(-/-)^* tadpoles (Suppl. Fig. 1C and D). The accumulation of autophagic shapes at the presynaptic terminal level, did not affect the density of synaptic vesicles or active zones, as they respectively remained at 106±15 and 1.75±0.4 in *LAMP2 ^(-/-)^* tadpoles (unpaired Student’s t-test, p=0.42 and p=0.31). The presence of a comparable synaptic vesicle pool organization could explain why neurotransmission was maintained in the neurons studied (Suppl. Fig. 1A). These results suggested that the overall function of retinotectal synapses was preserved in conditions of *LAMP2* deficiency or absence, and consequently, the deficits in visual responses were caused by retinal dysfunction rather than by alterations to central pathways.

Synapse specific presence of autophagic shapes in *LAMP2* mutant *X. tropicalis* tadpoles.

To further investigate how the lack of functional *LAMP2* expression affected the processing of sensory information, we next evaluated the accumulation of autophagic intermediates in synapses established by photoreceptors and olfactory sensory neurons (OSNs). The synaptic connections involving these two neuronal types share a functional similarity, as they create the first relay in vision or olfaction, but are morphologically distinct. Photoreceptors form ribbon synapses that tonically release glutamate following graded changes of the membrane potential. In contrast, OSNs establish conventional synapses that release glutamate upon the arrival of action potentials.

Rods established most of the contacts of photoreceptors with bipolar and horizontal neurons in the outer plexiform layer of the retina of WT *X. tropicalis* tadpoles (Fig. 7A). Presynaptic terminals contained a density of ∼500 cytoplasmic vesicles/μm^2^ and ribbon length was of 492±10 nm (n=29). Both parameters remained unaltered in *LAMP2 ^(-/-)^* tadpoles (Figs. 7B-G). In WT tadpoles, early endosomes covered 0.74% of the total surface inspected and were the most abundant element of the endolysosomal system (Fig. 7H). This observation is well aligned with previous studies carried out in ribbon synapses (Paillart et al. 2003) and suggested that synaptic vesicles underwent constitutive cycling supported by early endosomes. In *LAMP2 ^(-/-)^* tadpoles the presynaptic terminal surface covered by early endosomes remained at a similar value of 0.95% (Figs. 7A and B), however, there was a remarkable increase in autophagic shapes (Figs. 7C-F). In WT tadpoles, rod photoreceptor terminals were virtually devoid of autophagic intermediates but in *LAMP2* ^-/-^ tadpoles, they covered 2.28% of the surface inspected. Specifically, the presence of phagophores and autophagosomes (Figs. 7C and D) became obvious, late endosomes were more commonly observed (Fig. 7E) and some organelles related to degradation of intracellular elements appeared, such as multilamellar bodies (Fig. 7F). These results illustrate that autophagy normally plays a minor role in photoreceptor ribbon synapses, but nonetheless, it was impaired by the absence of *LAMP2*.

**Figure 7.**
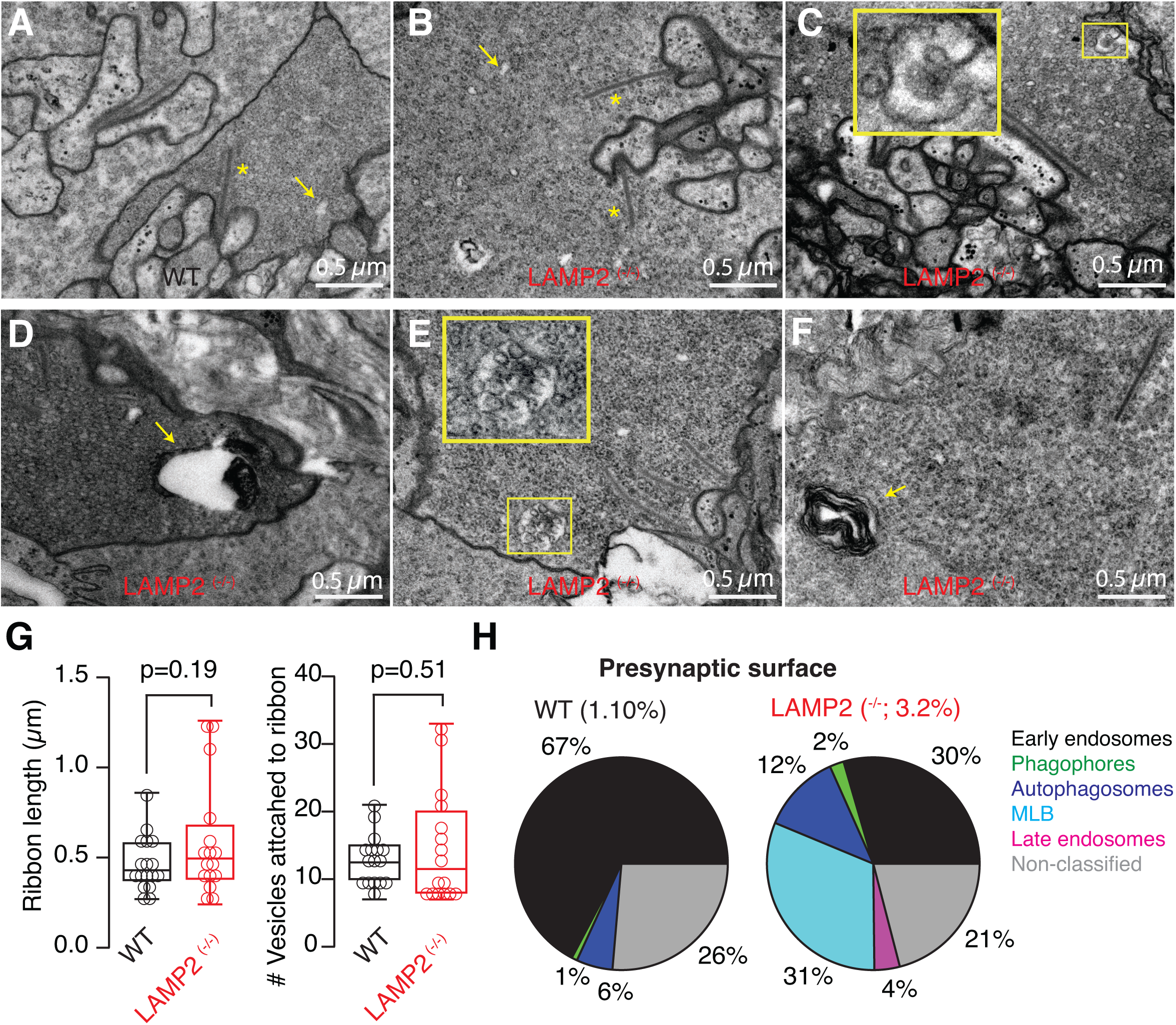
The presence of autophagic shapes is increased in photoreceptor ribbon synapses of *LAMP2* ^(-/-)^ tadpoles. **A)** Presynaptic terminal of a rod photoreceptor. Notice the presence of a high density of synaptic vesicles, ribbons (*) and early endosomes (arrow). **B)** *LAMP2* ^(-/-)^ showed photoreceptor ribbon synapses with a normal morphology containing ribbons and early endosomes. **C-F)** Presynaptic organelles related to the endolysosomal system and autophagic flux were found increased in *LAMP2* ^(-/-)^ tadpoles. Examples of a phagophore (C ), autophagosome (D), late endosome (E ) and a multilamellar body (F). **G)** Box plots showing the median (horizontal line), 25 to 75% quartiles (boxes), and ranges (whiskers) of ribbon length and the number of vesicles attached to ribbons in presynaptic terminals of photoreceptors found in WT and *LAMP2* ^(-/-)^ tadpoles. **H)** The surface of ribbon synapses covered by autophagic shapes and endolysosomal organelles was of 1.1% and 3.2% in WT and *LAMP2* ^(-/-)^ tadpoles, respectively. Pie charts illustrate the distribution of each type of organelle. MLB. Multilamellar bodies.

OSN axons enter the olfactory bulb and form the presynaptic element of glomeruli, which are postsynaptically defined by the dendrites of mitral/tufted cells. In WT tadpoles, the distinctive morphology of OSN axon terminals was characterized by the presence of a density of 215±1 synaptic vesicles/μm^2^ and 1.94±0.03 active zones/μm^2^ (Fig. 8A). In contrast, in *LAMP2 ^(-/-)^* tadpoles, values were reduced to 159±1 vesicles/μm^2^ and 0.22±0.01 active zones/μm^2^ (p<0.001, unpaired Student’s t-test). These alterations could be related to a likely dependency on autophagy to support normal presynaptic terminal function. Autophagic intermediates already occupied 2 % of axon terminal surface in WT tadpoles. Phagophores, autophagosomes and autolysosomes were commonly observed in OSN axon terminals (Fig. 8B). In *LAMP2 ^(-/-)^* tadpoles, there was an overall increase in autophagic shapes to cover 7.3% of the presynaptic surface inspected (Figs. 8C-G). The most remarkable change was related to phagophores (Fig. 8C) and autophagosomes (Fig. 8D), which increased by almost one order of magnitude. Organelles involved in degradative processes were also obvious (Fig. 8E), as well as mitochondria with an electrolucent matrix (Fig. 8F), which resembled those found in muscle fibers (Fig. 2C) and rods (Fig. 4G). Together, these observations suggested that endolysosomal function, autophagy and mitophagy were affected in OSN axon terminals.

**Figure 8.**
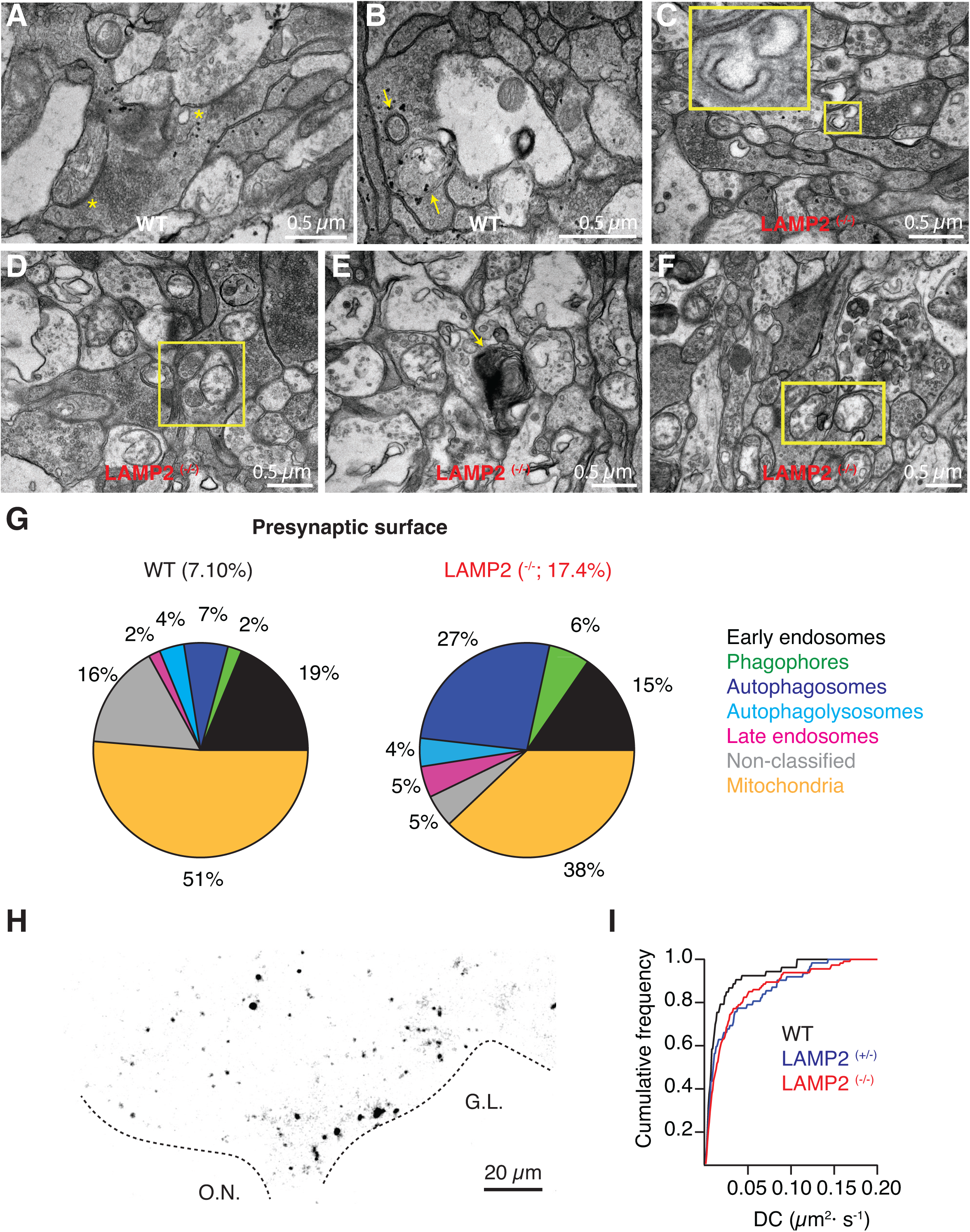
Increased presence of autophagic and endolysosomal organelles in presynaptic terminals of olfactory sensory neurons. **A)** Axon terminals of olfactory sensory neurons (OSNs) contain a high density of synaptic vesicles and multiple active zones (*) that primarily contact dendrites of mitral cells. **B)** Organelles of the autophagic flux such as autophagosomes are commonly observed in OSN presynaptic terminals of WT tadpoles. **C-E)** Autophagic intermediates are abundantly found in OSN presynaptic terminals of LAMP2 ^(-/-)^ tadpoles. Examples of a phagophore (C ), autophagosomes (D) and an autophagolysosome. **F)** Electrolucent mitochondria are present in OSN presynaptic terminals of LAMP2 ^(-/-)^ tadpoles.

Morphological observations were complemented by live imaging of acidic organelles in OSNs using lysotracker deep red (Fig. 8H). Previous observations showed that the mobility of endolysosomes present in OSN axon terminals increases upon synapse elimination (Terni and Llobet 2021). We found that the typical endolysosomal diffusion coefficients ranging from 0.05 to 0.2 μm^2^·s^-1^ showed an overall 40% enhancement in tadpoles lacking or deficient for *LAMP2* (Fig. 8I). This increase in mobility could be attributed to alterations in the cytoskeleton restricting organelle diffusion, and/or a shift towards a more mobile population of acidic organelles. Both conditions are consistent with the significant morphological alterations found in OSN axon terminals that can be summarized in a decrease in synaptic vesicles and an overall alteration of organelles related to the endolysosomal system and autophagy (Fig. 8). These findings suggest that the transfer of information could be altered in the olfactory bulb, thus being a novel trait of nervous system defects related to a *LAMP2* deficiency.

## DISCUSSION

The current study illustrates the unique potential of *X. tropicalis* to experimentally model human diseases. The knockout of *LAMP2* gene in *X. tropicalis* tadpoles replicates three typical alterations present in DD patients: skeletal muscle myopathy, cardiac alterations and visual defects. Comparable deficiencies have also been observed in mice (Saftig et al. 2001) and rats (Ma et al. 2018), which supports an evolutionary conserved role of LAMP2 protein in vertebrates (Qiao et al. 2023) despite a broken synteny. In mammals, the *LAMP2* gene is in the X chromosome and consequently, males are more severely affected than females, while in *X. tropicalis* it is in chromosome 8, an autosome. Here, we found similar changes in heterozygous (^+/-^) and homozygous (^-/-^) LAMP2 *X. tropicalis* in terms of mobility as well as visual and cardiac performance, which suggest the presence of a haploinsufficiency leading to characteristic dysfunctions present in DD. On this basis and considering that natural mating of LAMP2 (^-/-^ and ^+/-^) adult *X. tropicalis* generate >300 viable embryos, the experimental possibilities offered by this animal model system to study DD are exceptional. Two-week-old tadpoles (NF stage ∼50) are transparent and their behavior can be investigated using a variety of studies (Pratt and Khakhalin 2013) besides a wealth of available biochemical and physiological approaches (Tandon et al. 2017). For example, an improvement of mobility or cardiac deficits could be used to identify candidate drugs capable of ameliorating alterations present in DD patients. A goal that could be achieved by investigating swimming or cardiac performance in screens of compound libraries. This type of middle-throughput approach has been useful to detect two serotonin signaling molecules that could ameliorate gastrointestinal distress associated with autism (McCluskey et al. 2025), and, altogether evidence the potential of *X. tropicalis* as an experimental platform to investigate human diseases.

*LAMP2* deficient *X. tropicalis* tadpoles show significant visual deficits of retinal origin. Rods, which mediate green vision in *Xenopus* tadpoles(Röhlich and Szél 2000), are particularly affected. The alterations are obvious in mitochondria, as they contain an electrolucent matrix and a reduced number of cristae. This characteristic appearance of mitochondria is also observed in in skeletal muscle fibers and is likely related to the role played by LAMP2 in mitophagy (Hashem et al. 2017). In contrast, mitochondria found in cones show a normal appearance, which could explain the absence of deficits in the detection of red light (Röhlich and Szél 2000). The different perturbations to mitochondria found in rods and cones suggest that the role of LAMP2 must be interpreted in a cell-specific context. The individual contribution of each one of the three LAMP2 isoforms, LAMP2A, LAMP2B and LAMP2C, could be key as they show a variable distribution as well as different roles in autophagy and mitophagy (Kaushik et al. 2011; Qiao et al. 2023).

The cellular heterogeneity of LAMP2 functions could be an important determinant of the complex psychiatric alterations present in DD patients and, here, we provide evidence for a synapse-specific action. The presynaptic terminals of photoreceptors and OSNs define the first relay of vision and olfaction. But, despite this similarity related to the processing of sensory information, they are completely different in morphological and functional terms.

The transfer of information from photoreceptors to bipolar cells occurs in ribbon synapses (Lagnado and Schmitz 2015), whilst OSNs establish conventional synapses with mitral cells in olfactory glomeruli (Shepherd, Chen, and Greer 2004). The morphological analysis of both types of presynaptic terminals in WT tadpoles evidenced the normal presence of organelles of the endolysosomal system, such as early or late endosomes, in agreement with observations obtained in other species (Paillart et al. 2003; Kasowski, Kim, and Greer 1999). Autophagic intermediates, for example phagophores, were easily identified in OSN terminals but were absent from photoreceptor ribbon synapses. This difference illustrates that synapses rely on autophagy to perform their normal function (Vijayan and Verstreken 2017) but to a different extent. Endolysosomal and autophagic organelles increased by approximately threefold in the absence of *LAMP2* in both types of synapses, thus becoming particularly evident in conventional synapses established by OSNs. This finding cannot be extended to all synapse classes, because presynaptic terminals in the tectum did not show such a strong dependency on the endolysosomal system or autophagic flux. Our results are consistent with a synapse specific role of autophagy, likely determined or influenced by *LAMP2* isoforms. The need of synapses to maintain different levels of activity, tightly coupled to synaptic vesicle cycling, might be relevant to define the involvement of *LAMP2*, and consequently, the specific effects on neuronal circuits in its absence.

Several questions arising from these observations could be further explored. For example, the different contributions of LAMP2 isoforms to autophagy in the different synapses examined remains unknown. LAMP2A mediates chaperone-mediated autophagy, LAMP2B is involved in macroautophagy and LAMP2C participates in selective RNA and DNA degradation (Qiao et al. 2023). Their variable expression across neuronal types could underlie the synapse specific effects and isoform-specific rescue experiments in *X. tropicalis* could be a natural next step. Secondly, mTOR inhibition has been proposed as a possible therapy for heart deficits in DD, based on experiments in a zebrafish model (Dvornikov et al. 2019). This stems from the finding that impaired autophagic flux in LAMP2 mutants leads to a compensatory upregulation of mTOR signaling, this further suppresses autophagy initiation contributing to cellular damage. mTOR inhibitors would relieve this block, restoring autophagic flux through LAMP2-independent mechanisms and improving cardiac function. However, this might have the opposite effects in synapses where the accumulation of autophagic intermediates is already overwhelming the more constrained presynaptic compartments: mTOR inhibition might make this accumulation worse and lead to further synaptic dysfunction. Evaluating this possibility in *X. tropicalis* would be straightforward given the behavioral and electrophysiological readouts established in this study. Finally, the accumulation of autophagic intermediates in OSN terminals and the decrease in presynaptic vesicles, suggests that olfactory function may be compromised in DD. Olfactory dysfunction, to our knowledge, has not been formally addressed in patients with DD, and could be an unrecognized clinical feature of DD. Examining this question both experimentally using olfactory assays in *X. tropicalis,* as well as clinically, would further validate the translational aspect of this model and broaden our understanding of the neurological spectrum of DD.

## ACKNOWLEDGMENTS

Grant PID2021-124536NB-I00 (A.L.) from the Ministry of Science, Innovation and Universities (MICIU/AEI), co-funded by the European Regional Development Fund (ERDF), “a way of making Europe” supported work related to histological procedures and tadpole mobility assays. National Institute of Health (NIH) grant 2R25NS063307-41 (Marine Biological Laboratory, Neurobiology Course) supported electrophysiology, visual behavior and heart function work. NIH grants R24OD030008 (M.E.H.) and P40OD010997 (M.E.H.) supported generation and housing of mutant *X. tropicalis* lines. Tasks involved in this work were also supported by two Whitman fellowships (A.L.) in 2023 and 2024 (Marine Biological Laboratory, University of Chicago). The authors thank the institutional support from the María de Maeztu Unit of Excellence, Institute of Neurosciences, University of Barcelona, CEX2021-001159-M (Ministry of Science, Innovation and Universities) and the CERCA Program of Generalitat de Catalunya. A.L. is a Serra Húnter fellow.

