## Supplementary Figure 1 for "Synapse specific alterations of autophagy are a hallmark of Danon disease"

**SUPPLEMENTARY INFORMATION**

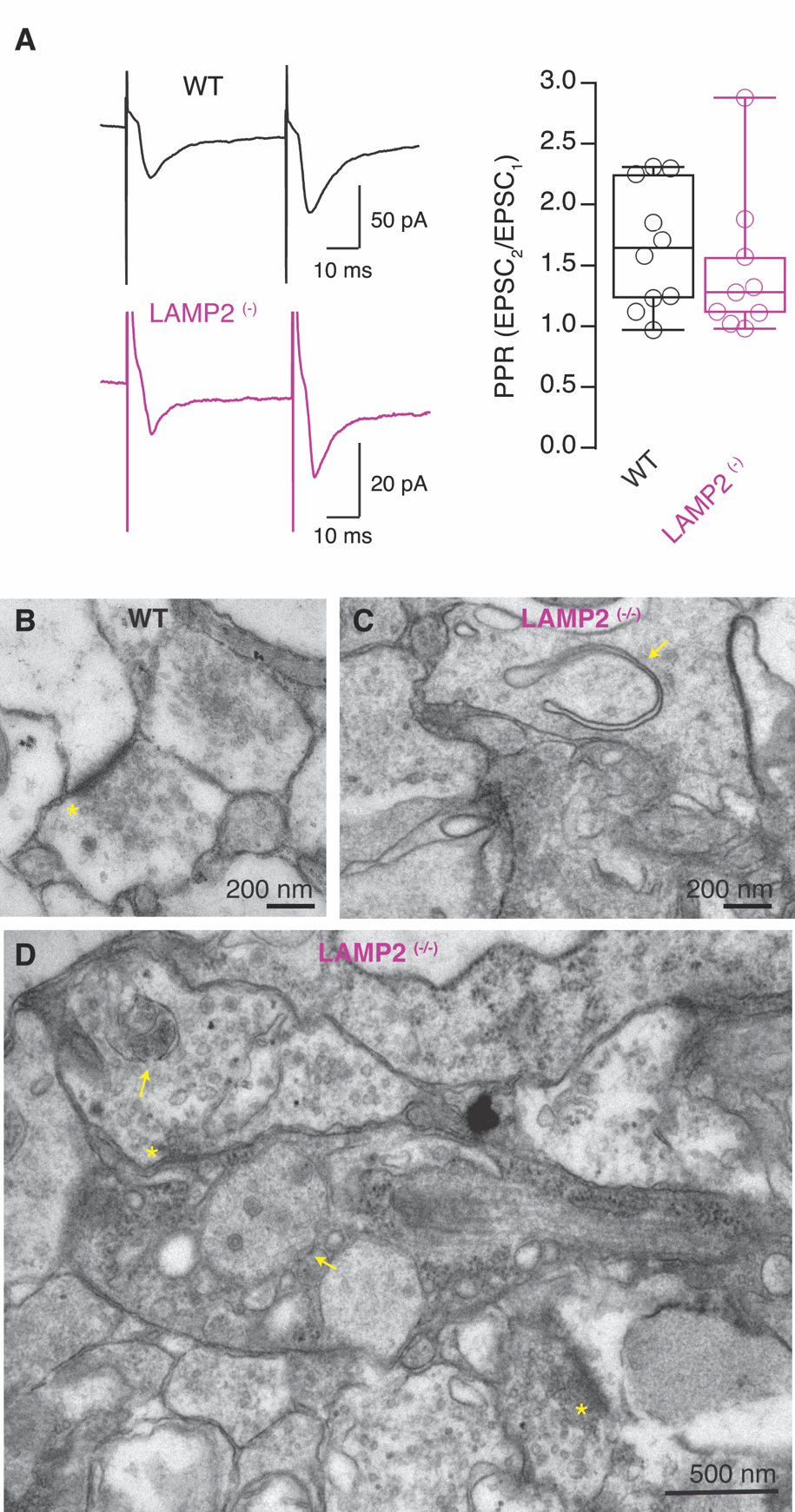

**Supplementary Figure 1. Centrally-evoked retinotectal activity remains normal in *LAMP2* ^(-)^ mutants. A)** Sample paired EPSCs recorded in tectal neurons, evoked by direct optic nerve stimulation, which bypasses the retina. Paired pulse facilitation, a measure of synaptic release competency remains unaltered in LAMP2 mutants, compared to WT tadpoles. **B)** Image of a conventional tectal synapse from a WT tadpole. **C,D)** Synapses found in the tectum of LAMP2 (-/-) tadpoles. Phagophores (C, arrow), were commonly observed. Autophagosomes were also obvious in pre and postsynaptic terminals (D, arrows). Asterisks indicate presynaptic active zones.
